# Cytonuclear discordance in the crowned-sparrows, *Zonotrichia atricapilla* and *Zonotrichia leucophrys*: a mitochondrial selective sweep?

**DOI:** 10.1101/2020.08.20.259549

**Authors:** Rebecca S. Taylor, Ashley C. Bramwell, Rute Clemente-Carvalho, Nicholas A. Cairns, Frances Bonier, Stephen C. Lougheed

## Abstract

The golden-crowned (*Zonotrichia atricapilla*) and white-crowned (*Z. leucophrys*) sparrows have been presented as a compelling case for rapid speciation. They display divergence in song and plumage with overlap in their breeding ranges implying reproductive isolation, but have almost identical mitochondrial genomes. Previous research proposed hybridization and subsequent mitochondrial introgression as an alternate explanation, but lacked robust nuclear gene trees to distinguish between introgression and incomplete lineage sorting. We test for signatures of these processes between *Z. atricapilla* and *Z. leucophrys*, and investigate the relationships among *Z. leucophrys* subspecies, using mitochondrial sequencing and a reduced representation nuclear genomic dataset. Contrary to the paraphyly evident in mitochondrial gene trees, we confirmed the reciprocal monophyly of *Z. atricapilla* and *Z. leucophrys* using large panels of single nucleotide polymorphism (SNPs). The pattern of cytonuclear discordance is consistent with limited, historical hybridization and mitochondrial introgression, rather than a recent origin and incomplete lineage sorting between recent sister species. We found evidence of nuclear phylogeographic structure within *Z. leucophrys* with two distinct clades. Altogether, our results support the true species status of *Z. atricapilla* and *Z. leucophrys*, and indicate deeper divergences between the two species than inferred using mitochondrial markers. Our results demonstrate the limitations of relying solely on mitochondrial DNA for taxonomy, and raise questions about the possibility of selection on the mitochondrial genome during temperature oscillations (e.g. during the Pleistocene). Historical mitochondrial introgression facilitated by past environmental changes could cause erroneous dating of lineage splitting in other taxa when based on mitochondrial DNA alone.

## 1. Introduction

Speciation is typically construed as an incremental process that requires millions of years to complete. For example, in sister species of birds, divergence in allopatry followed by secondary sympatry where the two taxa remain reproductively isolated despite possibilities for interbreeding, typically requires separation of at least 1.7 million years (Weir and Price, 2011). Against this backdrop, examples of very rapid speciation (hundreds of thousands rather than millions of years) can help to illuminate the ecological and genetic factors that underpin diversification and the evolution of reproductive isolation. There are various examples of purported rapid speciation in birds including goldfinches (*Carduelis*; Arnaiz-Villena et al., 1998), canaries (*Serinus*; Arnaiz-Villena et al., 1999), Darwin’s finches (Geospizinae; Grant, 1999), red crossbills (*Loxia curvirostra* complex; Benkman, 2003), indigobirds (*Vidua*; Sorenson et al., 2003), pheasants (Shen et al., 2010), and the southern capuchino seedeaters (*Sporophila*; Campagna et al., 2012). Rapid speciation in birds is typically accompanied by well-developed prezygotic isolating mechanisms (e.g. based on plumage, songs), despite sister species being genetically similar across most of their genome (Edwards et al., 2005). For example, the southern capuchinos comprise nine broadly sympatric species found in grasslands of South America that exhibit marked differentiation in male plumage and song, yet show virtually no molecular divergence, implying very recent origins within the second half of the Pleistocene (Campagna et al., 2012). Genomic evidence suggests that this remarkable diversification of male plumage is underlain by selection on regulatory elements of genes involved in melanogenesis (Campagna et al., 2017). The rapidity of divergence from a common ancestor in all of these avian systems means that intrinsic postzygotic isolation may evolve relatively more slowly making hybridization possible, and that prezygotic isolation may arise before we see complete reciprocal monophyly.

Golden-crowned (*Zonotrichia atricapilla*) and white-crowned (*Z. leucophrys*) sparrows have been touted as an example of exceedingly rapid speciation, perhaps splitting as recently as 50,000 years ago (Johnson and Cicero, 2004). The evidence for this is not unequivocal. Early genealogies based on allozyme data (Zink, 1982) and mitochondrial DNA (mtDNA) (Zink et al., 1991) implied that *Z. atricapilla* and *Z. leucophrys* were independently-evolving sister species. However, subsequent analyses of mtDNA sequences suggested that they are not reciprocally monophyletic, even sharing some haplotypes (Weckstein et al., 2001; McKereghan-Dares, 2008;), and that their low global mitochondrial nucleotide diversity is comparable to a single, homogenous avian population rather than two well diverged species. Weckstein et al. (2001) presented alternate hypotheses to explain this pattern: 1) *Z. atricapilla* and *Z. leucophrys* originated in allopatry, followed by contact, historical hybridization and subsequent introgression, with ultimate replacement of an one ancestral mitochondrial genome in one of the species with that of the other (mtDNA capture; see Good et al., 2008); or 2) these two species originated very recently, with insufficient time for complete lineage sorting to have occurred.

The white-crowned sparrow has a geographic range spanning most of North America, while the golden-crowned is generally confined to regions west of the Rocky Mountains (Cortopassi and Mewaldt, 1965; Fig. 1). Both species are migratory, breeding in the northern parts of their ranges and overwintering in the south, although *Z. leucophrys* have some year-round resident populations along the coast of Washington, Oregon and California. Adults are distinct in their plumages, breeding behaviours, and vocalizations (del Hoyo et al., 2011). Playback experiments have shown that both sexes of *Z. leucophrys* react more strongly to conspecific than heterospecific songs (Pleszczynska, 1979), and to natal versus non-natal vocal dialects (Baker et al., 1982). Female *Z. l. nuttallii* mate assortatively with males of their natal dialect population (Tomback and Baker, 1984). Finally, while some reports of *Z. atricapilla-Z. leucophrys* hybrids exist (Miller, 1940; Morton and Mewaldt, 1960; McCarthy, 2006), these represent a comparatively small number given the extensive overlap in their breeding distributions across British Columbia, the Yukon, and Alaska (Cortopassi and Mewaldt, 1965). All existing evidence thus suggests that these are good biological species.

**Fig. 1.**
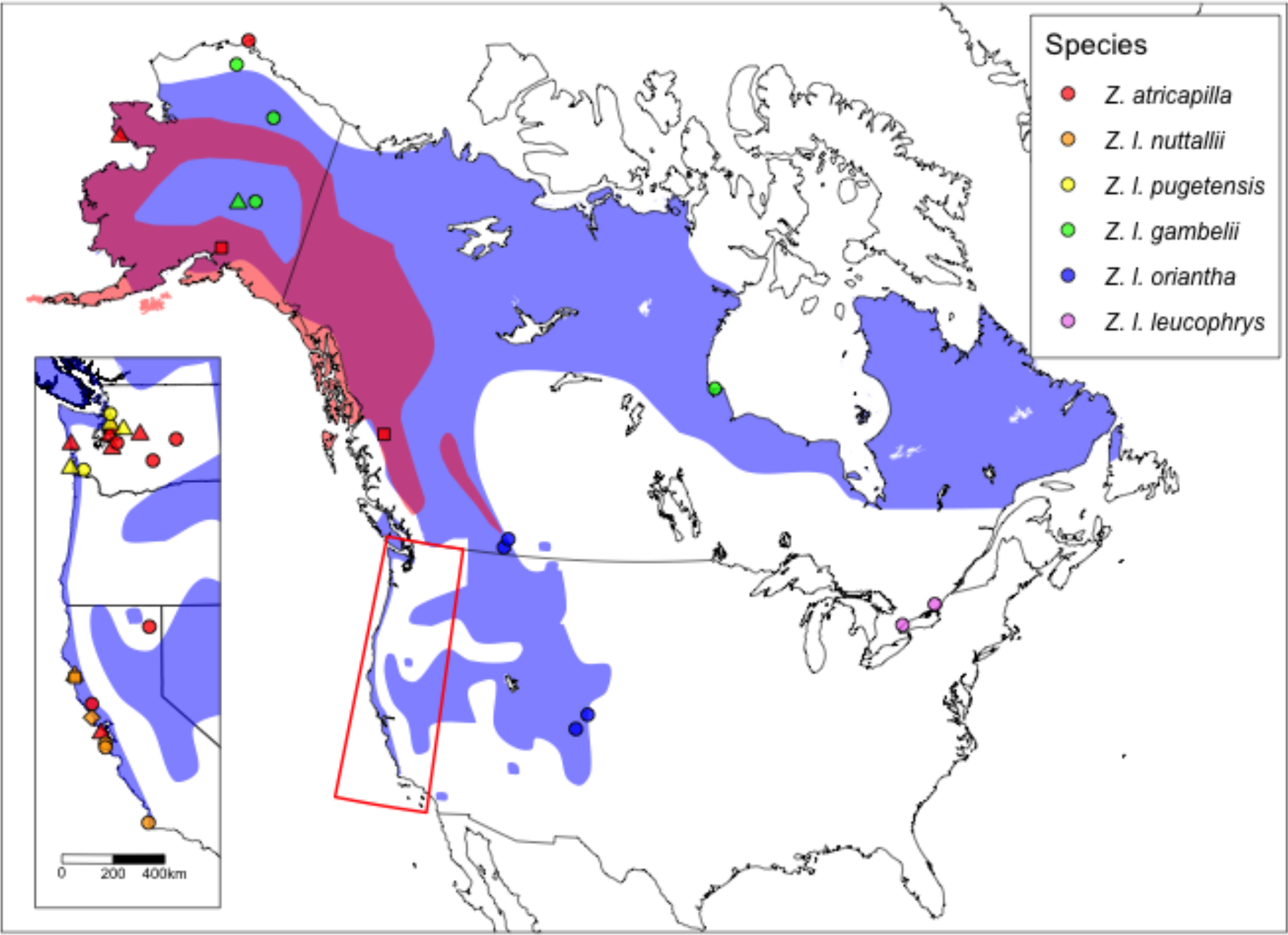
The locations of sampling sites of *Z. atricapilla* and *Z. leucophrys* individuals The background colors represent the breeding ranges, pink for *Z. atricapilla* and purple for *Z. leucophrys* with deep pink representing range overlap. Circles indicate samples used for both ddRADseq and mitochondrial DNA sequencing, triangles represent locations where at least one sample has mitochondrial DNA only, squares represent locations where at least one sample has ddRAD data only. More sampling details are provided in Table 1.

Within *Z. leucophrys* there are five named subspecies (*Z. l. nuttallii, Z. l. pugetensis, Z. l. gambelii, Z. l. oriantha* and *Z. l. leucophrys).* A recent genomic study of nuclear DNA investigated population structure across a contact zone of *Z. l. nuttallii*, and *Z. l. pugetensis.* Differentiation was found between the two subspecies, driven by differences in advertising song, indicating that they are separate evolutionary significant units (Lipshutz et al., 2017). However, secondary contact has led to ongoing hybridization between the two subspecies (Lipshutz et al., 2017; Toews, 2017). Genetic relationships based on nuclear DNA among the other subspecies are largely unknown.

The five crowned-sparrow species (golden-crowned and white-crowned sparrows along with Harris’ sparrow, *Z. querula*, white-throated sparrow, *Z. albicollis*, and the rufous-collared sparrow, *Z. capensis*) have contributed significantly to our current understanding of speciation with research on reproductive isolation (e.g. Danner et al., 2011), mating behaviour (e.g. Tomback and Baker, 1984; Houtman and Falls, 1994; Lipshutz et al., 2017), phenology (e.g. Handford, 1980, Moore et al., 2005), migration (e.g. Landys et al., 2004; McFarlan et al., 2009; Price et al., 2010), vocal dialects, song learning, and function (e.g. Handford and Nottebohm, 1976; Lougheed and Handford, 1992; MacDougall-Shackleton and MacDougall-Shackleton, 2001; Shizuka, 2014, Toews, 2017), and phylogeography, geographic range expansions, and secondary contact zones (Lougheed et al., 2013; Campagna et al., 2014). Despite their prominence in evolutionary and ecological studies, there is phylogenetic uncertainty about the relationships among the species (Carson and Spicer, 2003).

Here, we use a genome-wide panel of nuclear Single Nucleotide Polymorphisms (SNPs) combined with mitochondrial DNA sequencing to investigate relationships between *Z. atricapilla* and *Z. leucophrys* to test the alternate propositions presented by Weckstein et al. (2001). If the species originated, as in the first hypothesis, with allopatric divergence and the introgression of mitochondrial DNA from one species into another, we expect signatures of genetic differentiation and monophyly in the nuclear DNA. Conversely, under the second hypothesis with recent speciation we predict low genetic differentiation and a lack of reciprocal monophyly in the nuclear DNA. We include samples from all subspecies within *Z. leucophrys*, and from all five crowned sparrow species so that we can assess the phylogenetic affinities of these taxa.

## 2. Materials and methods

### 2.1. Sampling

From museum collections and our own sample archive, we assembled 73 tissue and blood samples of *Z. atricapilla* and *Z. leucophrys* from across their respective geographic ranges, including individuals from all five subspecies of *Z. leucophrys* (Fig. 1; Table 1) for ddRAD sequencing. For mtDNA sequencing, 74 samples were used (Table 1). We added eight specimens from the remaining three *Zonotrichia* species (two from *Z. albicollis*, three from *Z. capensis*, two from *Z. querula*; Table 1; Fig. S1). Tissue samples were preserved in lysis buffer (4.0M urea, 0.2M NaCl, 10mM EDTA, 0.5% n-Lauroyl Sarcosine, 100mM Tris-CHl; pH 8.0) or 99% ethanol until extraction.

**Table 1.**
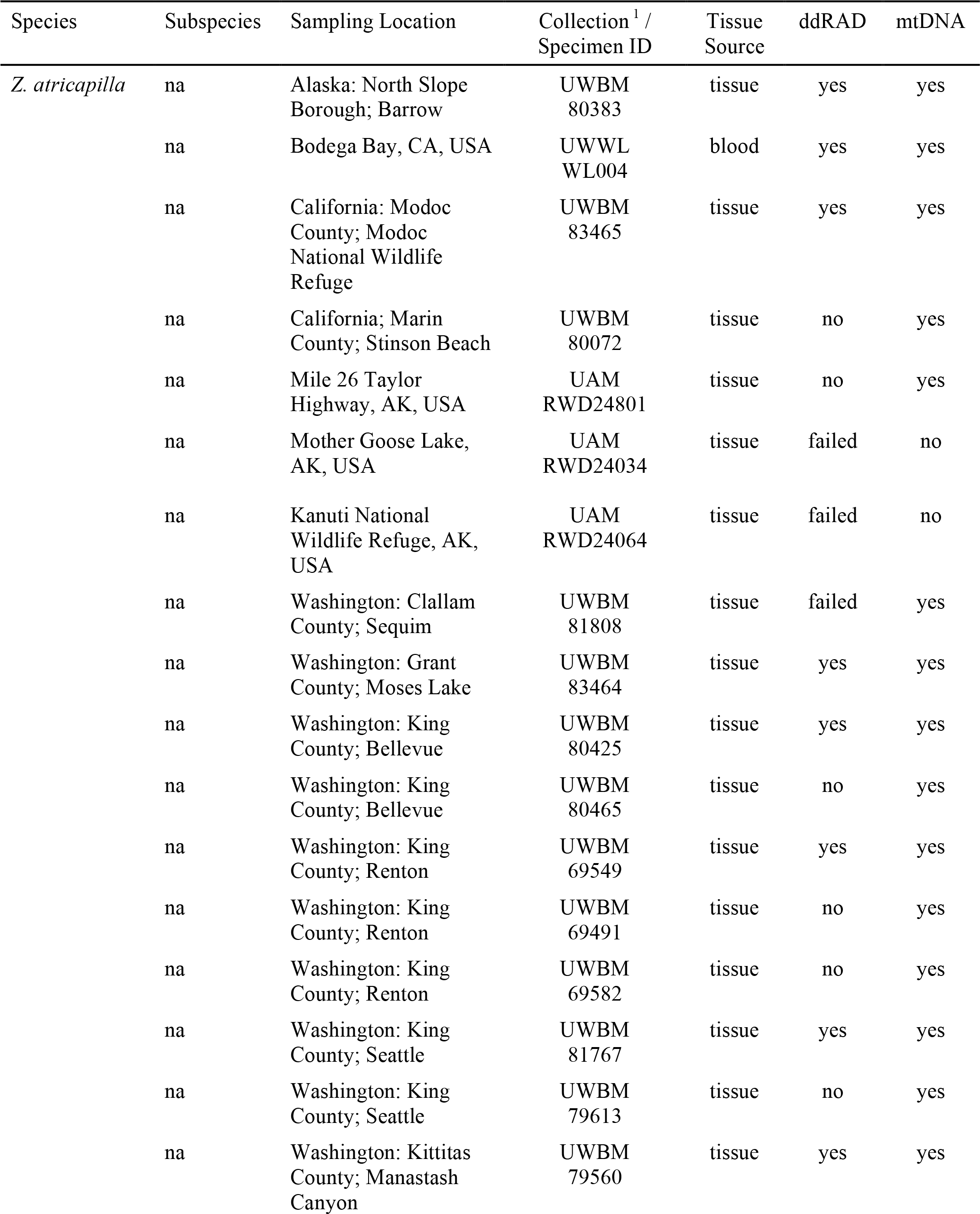
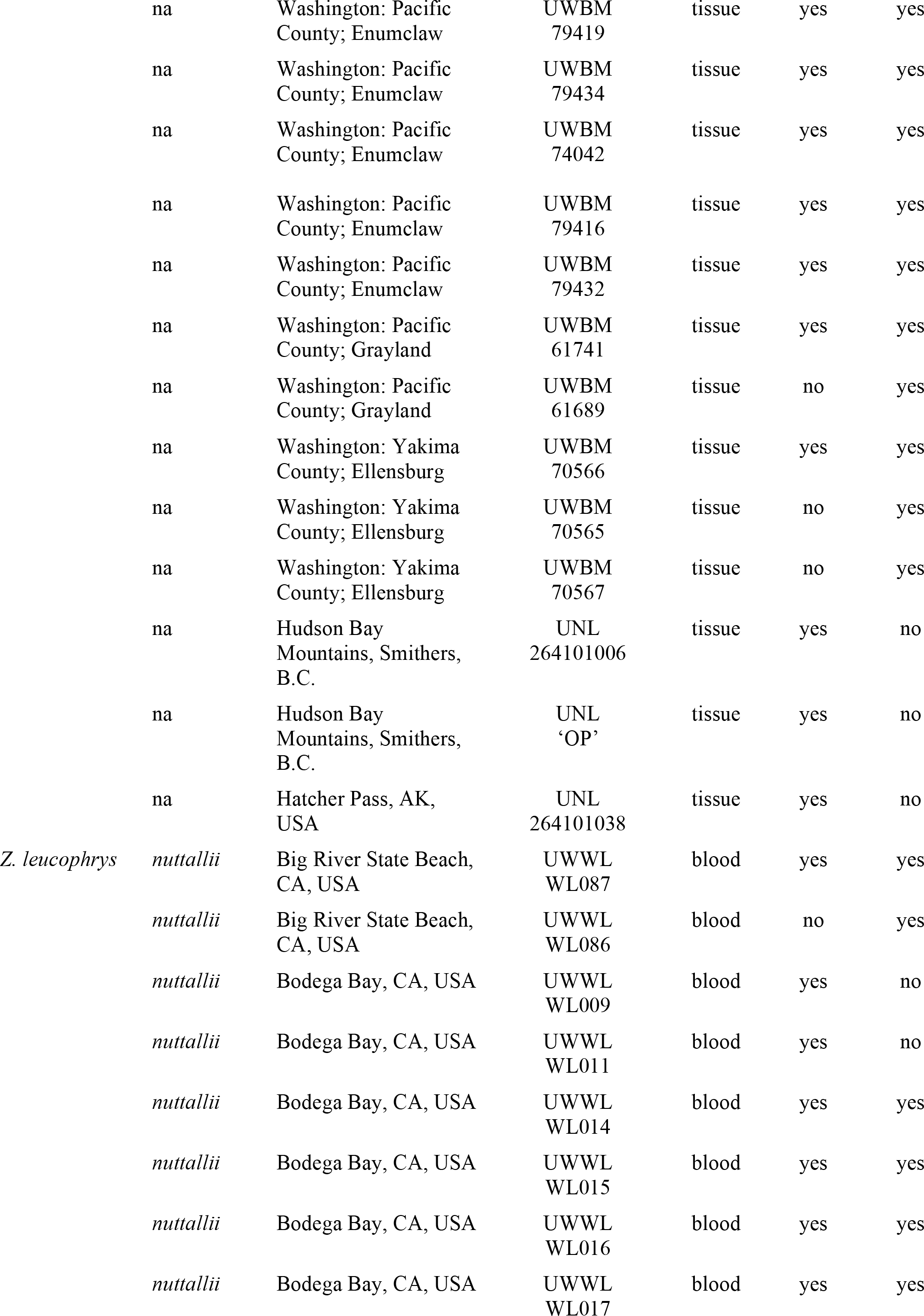
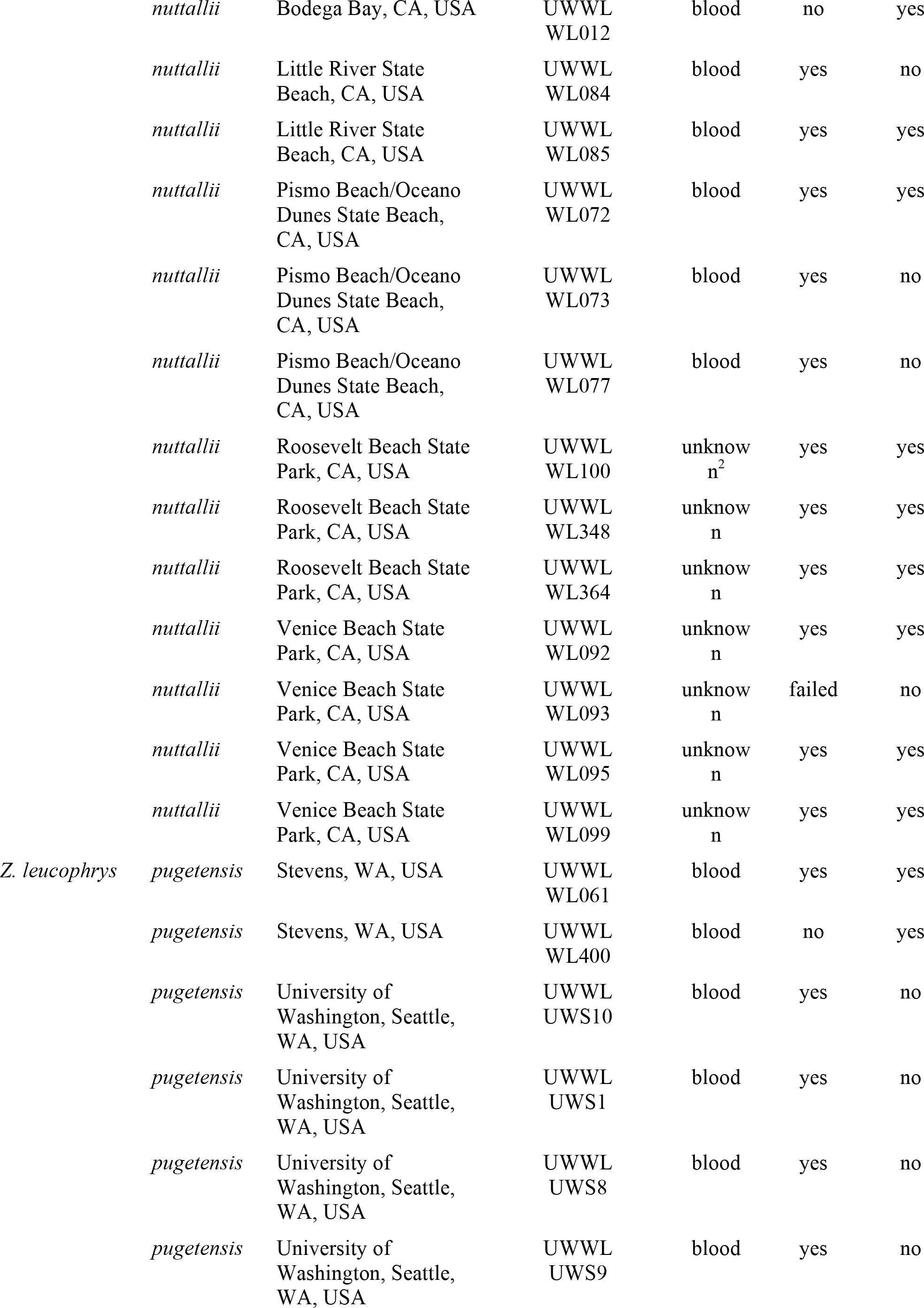
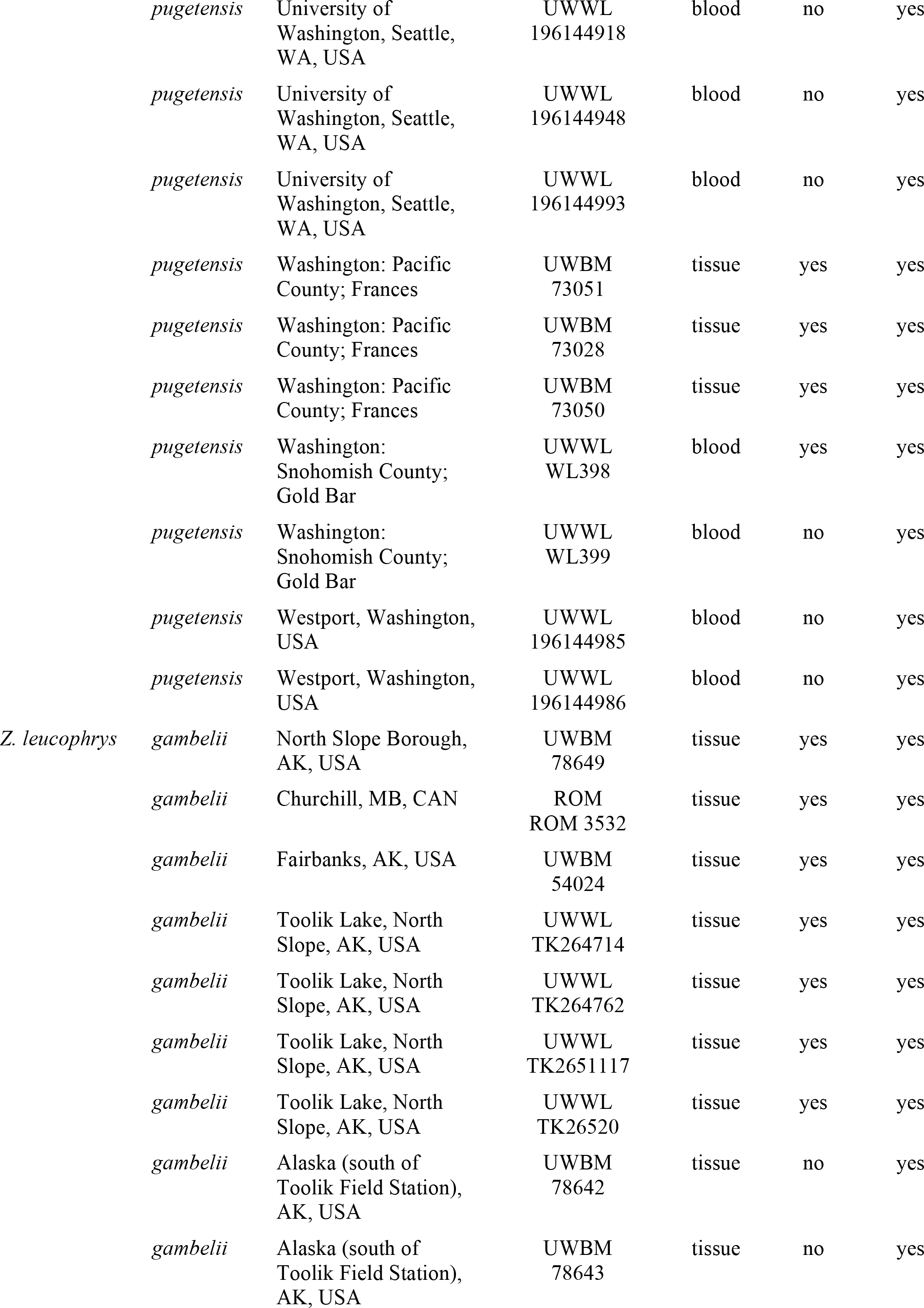
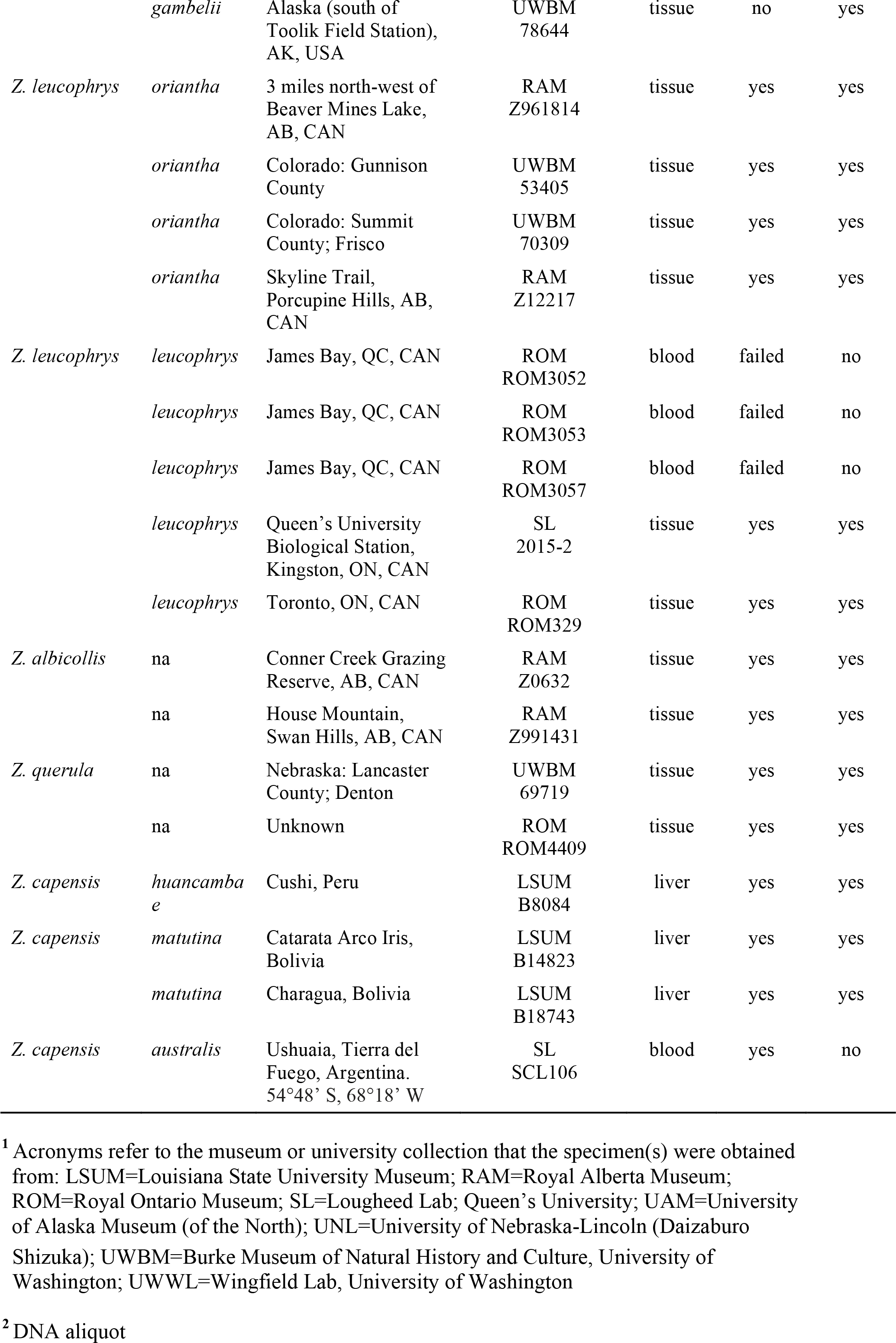
Species and subspecies designations, sampling locales, collection, tissue source, and datasets for specimens used in this study (ddRAD and mtDNA).

### 2.2. DNA extraction

Total genomic DNA was extracted either using the DNeasy^®^ Blood and Tissue Kit (QIAGEN, Mississauga, Ontario, Canada) following the manufacturer’s protocols, or a standard phenol-chloroform method (Sambrook & Russell, 2001). For ethanol preserved samples, excess ethanol was allowed to evaporate before extraction. Samples fixed in lysis buffer were washed in a 1x phosphate-buffered saline (PBS; 10mM, pH 7.4) solution. All newly extracted samples were treated with RNase A (10 mg/mL) and incubated at 37°C for 30 minutes. We checked the purity of our DNA using a Nanodrop ND-1000 spectrophotometer (Nanodrop Technologies Inc., Wilmington, DE, USA). If needed, due to low concentration or purity, samples were re-precipitated using a standard ethanol precipitation protocol with sodium acetate (3M, pH 5.2) as the carrier molecule, and then re-suspended in microfiltered autoclaved water and stored at −20°C until subsequent analyses. We quantified the concentration of double-stranded DNA in each sample using a Qubit® 2.0 Fluorometer (Life Technologies, Grand Island, NY, USA).

### 2.3. Mitochondrial DNA sequencing

Mitochondrial DNA sequence variation was assayed from control region (approximately 413bp) and cytochrome oxidase I (COI, 666bp) using the primers, PCR cocktail recipes, and cycling parameters given in Lougheed et al. (2013; primers: LCA1271-R and ZNGluF3) and Kerr et al. (2009; primers BirdF1 and COIbirdR2), respectively. The amplification products were visualized on 1.0% agarose gels using RedSafe™ Nucleic Acid Staining Solution, and purified using AMPure XP 2X Solid Phase Reversible Immobilization (SPRI) following the manufacturer’s instructions. Purified products were sent to The Centre for Applied Genomics, SickKids Hospital, Toronto, ON, Canada) for sequencing on an ABI 3730XL using the BigDye™ terminator kit. Mitochondrial sequences will be deposited in Genbank upon acceptance.

### 2.4. Double-digest restriction associated DNA (ddRAD) sequencing

Following Poland et al. (2012) double-digest RAD-Seq libraries were constructed at the Institute for Integrative Systems Biology (IBIS; at Laval University) using the restriction enzyme pairing of SbfI and MspI. Uniquely barcoded samples were pooled into a library and then size selected for roughly 100-300 bp using a BluePippin® DNA size selection system (Sage Science Inc., Beverly, MA, USA). The library was forwarded to Génome Québec (Génome Québec Innovation Centre) and assessed using a LabChip GX (Perkin-Elmer, MA, USA) instrument to determine library size quality. The library was then sequenced as single-end, 100 base-pair reads on a single Illumina HiSeq2000 lane (Illumina, San Diego, CA, USA). All ddRADseq data will be archived on the Dryad Digital Repository upon acceptance.

### 2.5. ddRADseq read filtering

Raw reads were de-multiplexed sequentially by barcode length using Stacks 2.3 (Catchen et al., 2013). Low quality reads (with a Phred score below 10), and those with uncalled bases, were filtered out and reads were truncated to 92bp. De-multiplexed reads were aligned to the *Zonotrichia albocolis* genome from NCBI (GCF000385455, Bioproject PRJNA197293) using the ‘very_sensitive’ setting in Bowtie2 (Langmead and Salzberg, 2012). Resulting SAM files were converted to BAM files and sorted using Samtools 1.1 (Li et al., 2009). We then used the Refmap and Populations modules in Stacks randomly selecting one SNP per locus, filtering out loci with a minor allele frequency of less than 0.05. After an initial run, seven individuals were found to have high levels of missing data and were removed, leaving 66 individuals (Table 1). We re-ran Refmap and Populations without the low-quality samples using two filtering settings, one allowing for 10% missing data (all individuals as one population and setting r to 0.9) and the other without any missing data (setting r to 1.0). We also did the same but without the outgroup taxa to create input files with only *Z. atricapilla* and *Z. leucophrys*, leaving 58 individuals (*18 Z. atricapilla*, 18 *Z. l. nuttalli*, 9 *Z. l. pugetensis*, 7 *Z. l. gambelli*, 4 *Z. l. oriantha*, and 2 *Z. l. leucophrys*). All analyses were conducted with both datasets (with no missing data and with 10% missing data).

### 2.6. mtDNA network and phylogenetics

We used the concatenated COI and control region sequences (total 1211 bp) to make a statistical parsimony haplotype network in PopArt (Clement et al., 2000; Leigh and Bryant, 2015; PopArt: http://popart.otago.ac.nz) using individuals from *Z. leucophrys* and *Z. atricapilla*. We performed Bayesian phylogenetic analyses on a dataset comprising all individuals and a dataset with only individual haplotypes identified in our network analysis. The best partitioning scheme and models of evolution for each dataset were selected using PartitionFinder2 (Lanfear et al., 2016). For the full dataset the best partition scheme and associated of models of evolution were: HKY+I for control region, HKY+I for COI codon position 1, F81 for COI codon position 2, and HKY+I for COI codon position 3. For the haplotype dataset the best partition scheme and associated of models of evolution were: HKY+I for control region, HKY+I for COI codon position 1, F81 for COI codon position 2, and HKY for COI codon position 3.

We inferred phylogenetic relationships using the full and haplotype mtDNA datasets using Bayesian inference with MrBayes v3.2.6 x64 (Huelsenbeck and Ronquist, 2001, Ronquist et al., 2012). For all models, partitions were unlinked for substitution rates and proportion of invariant sites (I). allowing parameter values to vary independently. Each Bayesian analyses included two simultaneous runs each with random starting trees, one cold and three incrementally heated Markov chains, and default priors for all parameters. Each analysis was allowed to run until the standard deviation of split frequencies < 0.01, sampling every 100 generations (full dataset: 2 million generations; haplotype dataset: 1 million generations). We visually assessed that our runs had achieved stationarity by evaluating the likelihoods in the program Tracer version 1.7.1 (Rambaut et al., 2018). For each analysis, we constructed a 50% majority rule consensus tree (discarding the first 25% of trees as burnin).

### 2.7. Phylogenomic reconstruction using SNPs

Using the PHYLIP files produced in Stacks, we ran RAxML version 8 (Stamatakis, 2014) to generate a maximum likelihood tree. We used a General Time Reversible model with a gamma distribution of substitution rates and a proportion of invariant sites (GTRGAMMAI), and used the rapid bootstrap algorithm using 1000 replicates. We ran the analysis will all species, but because of the long branch lengths (see Results), we also ran the analysis using *Z. querula* as an outgroup with only *Z. atricapilla* and *Z. leucophrys* individuals to make relationships clearer.

We ran Treemix 1.13 (Pickrell and Pritchard, 2012) to infer a maximum likelihood tree and to investigate any migration between *Z. atricapilla* and/or *Z. leucophyrs* sub-species. We ran Treemix from 0-9 migration events, running it without grouping SNPs for three iterations, and grouping SNPs into windows of 20 for three iterations to account for possible linkage and because the OptM package (below) needs different likelihood scored between iterations. We plotted the results in RStudio 1.0.136 (RStudio Team, 2015) and used the R package OptM (available here: https://cran.r-project.org/web/packages/OptM/index.html) to calculate the ad hoc statistic delta M, which is the second order rate of change in the log-likelihood of the different migration events, to help infer how many migration events to visualize.

### 2.8. Population structure

All population structure analyses were done using a dataset comprising only *Z. atricapilla* and *Z. leucophrys.* We ran an ADMIXTURE analysis (version 1.3; Alexander et al., 2009) for population assignment. To do this, we converted the VCF file into a BED file using Plink 1.9 (Purcell et al., 2007). We pruned the dataset to remove any SNPs with a correlation coefficient (R^2^) of 0.1 or greater. We then ran ADMIXTURE from K 1 to 8, and plotted the results in RStudio. We also did a principal component analysis in RStudio using the packages ADEGENET (Jombart, 2008) and vcfR (Knaus and Grünwald, 2017). We did an additional PCA on a subset of data with *Z. l. nuttalli* and *Z. l. pugetensis* only, and another PCA on *Z. l. gambelli*, *Z. l. oriantha*, and *Z. l. leucophrys* only to investigate possible substructure (see Results). We calculated pairwise Fst between *Z. atricapilla* and all *Z. leucophrys* subspecies, as well as between the three major lineages uncovered in the analyses (see Results) in RStudio using the HIERFSTAT package in R (Goudet, 2005) and the Weir and Cockerham calculation (WC84; Weir and Cockerham, 1984).

## 3. Results

### 3.1. ddRADseq data quality

We recovered 18.4 million single-end Illumina reads. After removing individuals with low quality reads, we had 3,469 SNPs in the VCF file with no missing data and 7,096 SNPs in the VCF file allowing 10% missing data for the datasets with all five crowned sparrow species, and 4,139 SNPs in the VCF file with no missing data and 6,626 SNPs in the VCF file allowing 10% missing data in the files with *Z atricapilla* and *Z. leucophrys* only. The depth was high for all retained individuals, ranging from 41.6 – 183.1X (aside from one *Z*. *capensis* individual with 481.4 X) for the file with outgroups and no missing data, and between 39.0 – 183.3X (aside from one *Z. capensis* individual which has 464.1 X) in the file allowing 10% missing data. These datasets were used for the RAxML analyses. Similarly, for the dataset comprising *Z atricapilla* and *Z. leucophrys* only the depth per individual ranged from 42.7 – 183.0X for the file with no missing data and between 39.7 – 169.3 X for the file allowing 10% missing data. These datasets were used for all subsequent nuclear SNP analyses, however all results presented are from the analyses with no missing data as all results were the same.

### 3.2. Evolutionary relationships apparent from analysis of mitochondrial DNA

The most obvious feature of both the mtDNA network (Fig. 2) and our Bayesian tree from analysis of distinct mtDNA haplotypes (Fig. 3) or of all individuals (Fig. S2) is that *Z. atricapilla* and *Z. leucophrys* share mtDNA haplotypes and do not form distinct, reciprocally monophyletic clusters/clades. We also find no obvious phylogeographic structure among *Z. leucophrys* subspecies, although some haplotypes are unique to particular subspecies. *Zonotrichia albicollis* and *Z. querula* comprise sister lineages, that are in turn, sister to the *Z. atricappila*/*Z. leucophrys* clade. The North American crowned sparrow clade is reciprocally monophyletic with the rufous-collared sparrow, *Z. capensis*.

**Fig. 2.**
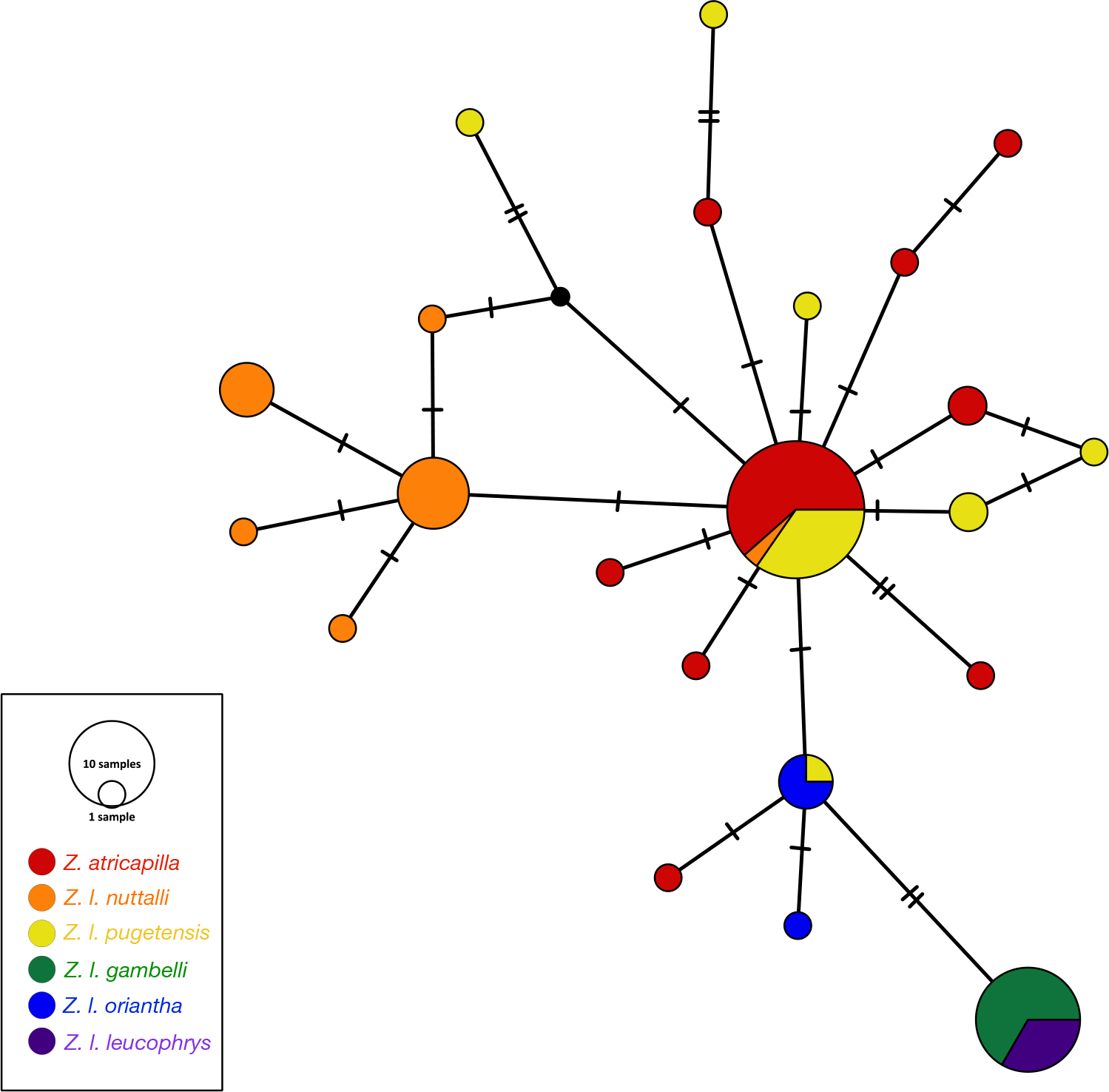
Statistical parsimony network of 1211 bp of the mitochondrial control region and COI concatenated for *Z. atricapilla* and all *Z. leucophrys* subspecies. Black circles represent unsampled or now extinct haplotypes and the hatch marks indicate mutations. Circle size indicates the number of samples of each haplotype.

**Fig. 3.**
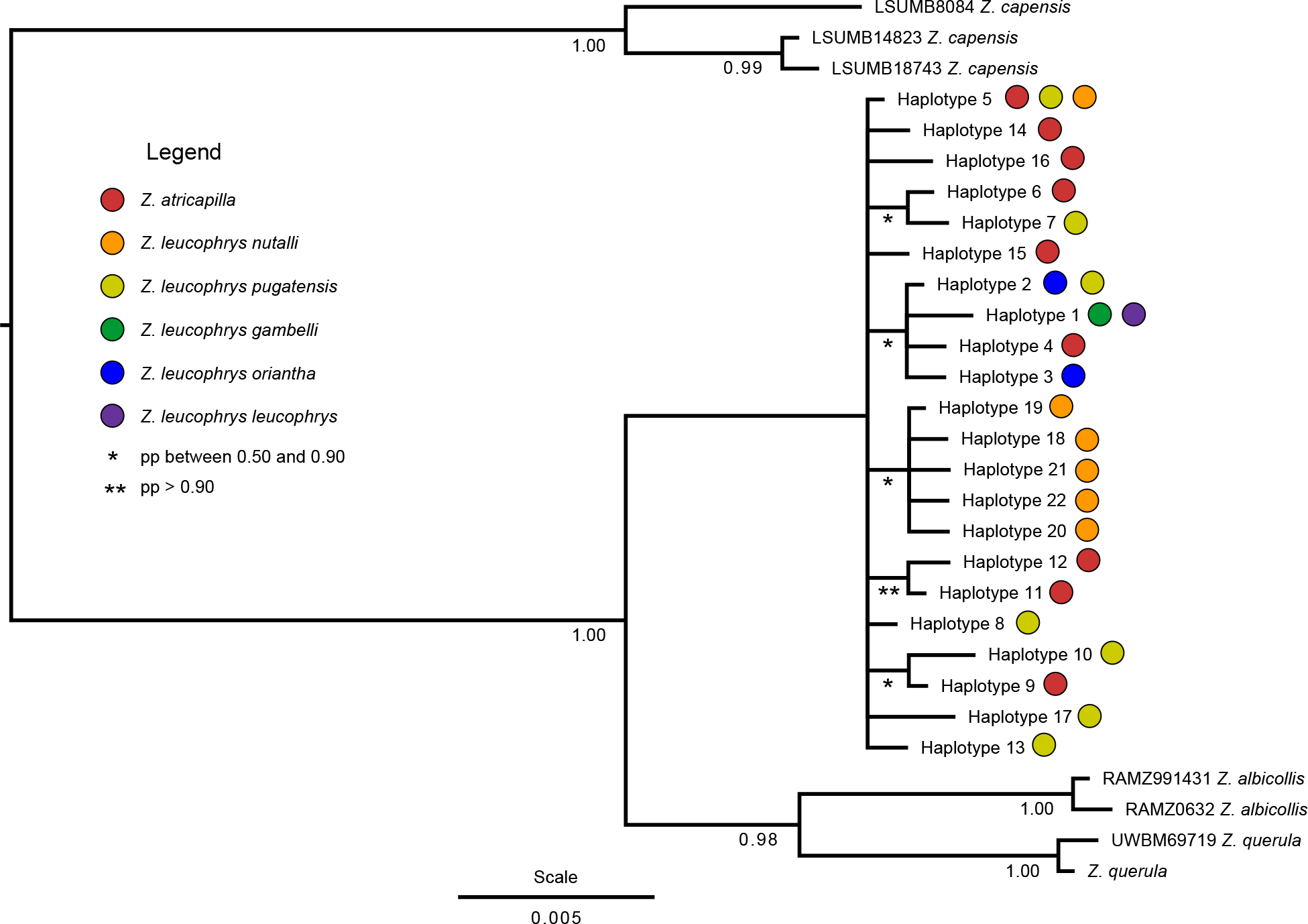
Bayesian mitochondrial DNA phylogenetic analyses using individual haplotypes from all five *Zonotrichia* subspecies. The legend shows which haplotypes are present in *Z. atricapilla* and the *Z. leucophrys* subspecies.

### 3.3. Interspecific relationships and population structure in Zonotrichia with nuclear DNA

In contrast to our results from analysis of mtDNA, *Z. atricapilla* and *Z. leucophrys* are well-supported, reciprocally monophyletic sister taxa in the RAxML phylogenomic analysis (Fig. 4A and 4B). The *Z. leucophrys* lineage was further divided into two clades: *Z. l. nuttallii* and *Z. l. pugetensis* (NP), and *Z. l. gambelii*, *Z. l. oriantha*, and *Z. l. leucophrys* (GOL). *Zonotrichia queula* was sister to *Z. atricapilla* and *Z. leucophrys, followed by Z. albicolis*, with *Z. capensis* recovered as sister to all North American congeners (Fig. 4A and 4B).

**Fig. 4.**
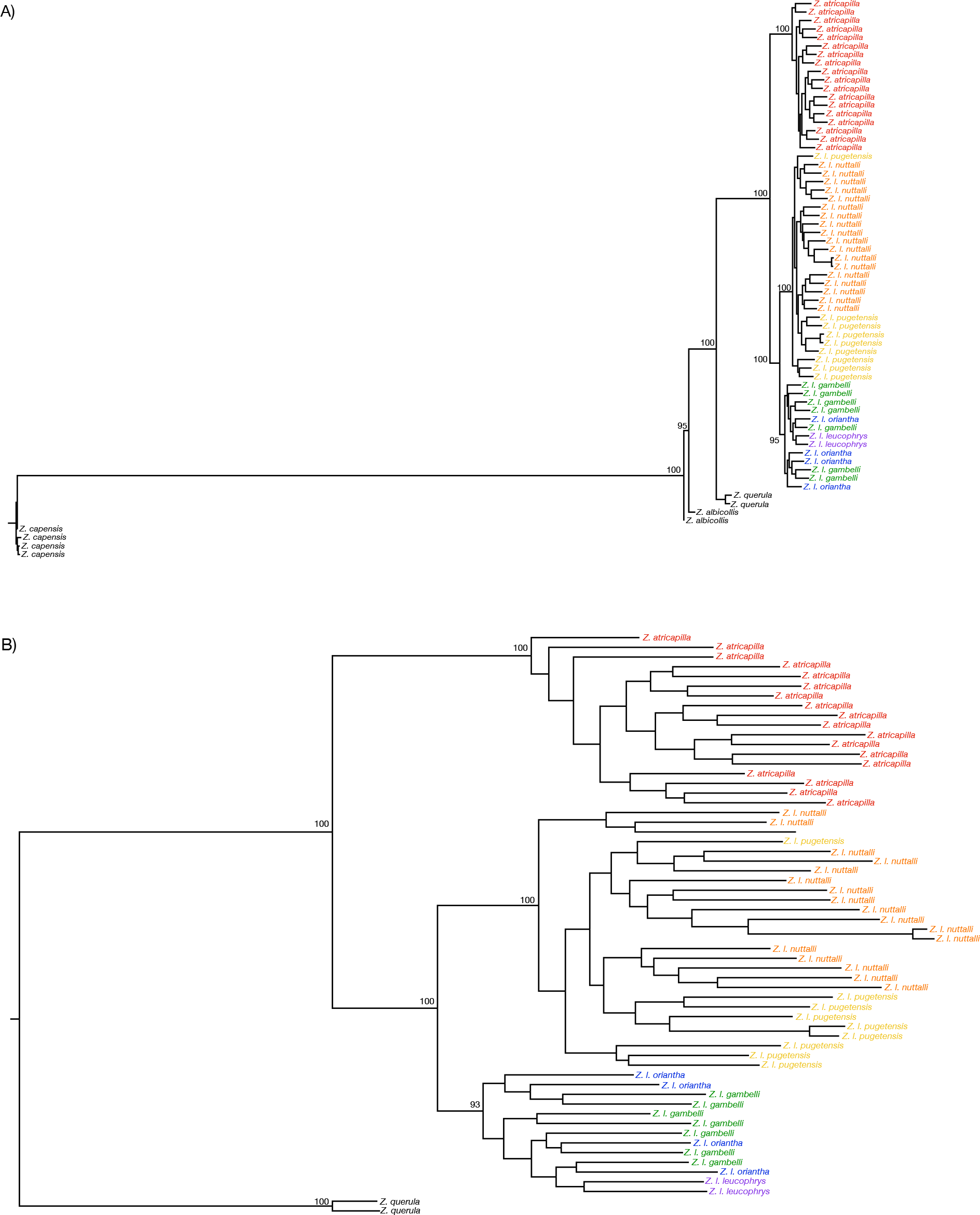
Maximum-likelihood trees produced in RAxML with all outgroups (A) and with only one outgroup taxa to show relationships within *Z. leucophrys* more clearly (B). Node labels indicate bootstrap support values from 1000 bootstrap replicates; bootstrap values above 90% are shown. Name labels are coloured according to species and subspecies (red=*Z. atricapilla*; orange=*Z. l. nuttallii*; yellow=*Z. l. pugetensis*; green=*Z. l. gambelii*; purple=*Z. l. leucophrys*; blue=*Z. l. oriantha*; black=outgroup taxa).

The Treemix maximum likelihood tree was similar, with *Z. atricapilla* well separated from all Z*. leucophrys* subspecies, and showing a large drift parameter (Fig. 5A). The NP clade remains distinct from the GOL subspecies, and shows evidence of some drift (Fig. 5A). Treemix stopped adding migration events after 5, and OptM indicated that one migration event which shows migration between the ancestor of the NP clade into *Z. l. leucophrys* (Fig. 5B; Fig S3). When adding three migration events, the analysis did infer migration from *Z. atricapilla* into *Z. l. leucophrys* (Fig. 5D); however adding three migration events did not increase the variance explained compared with adding one or two migration event by a lot (Fig S3).

**Fig. 5.**
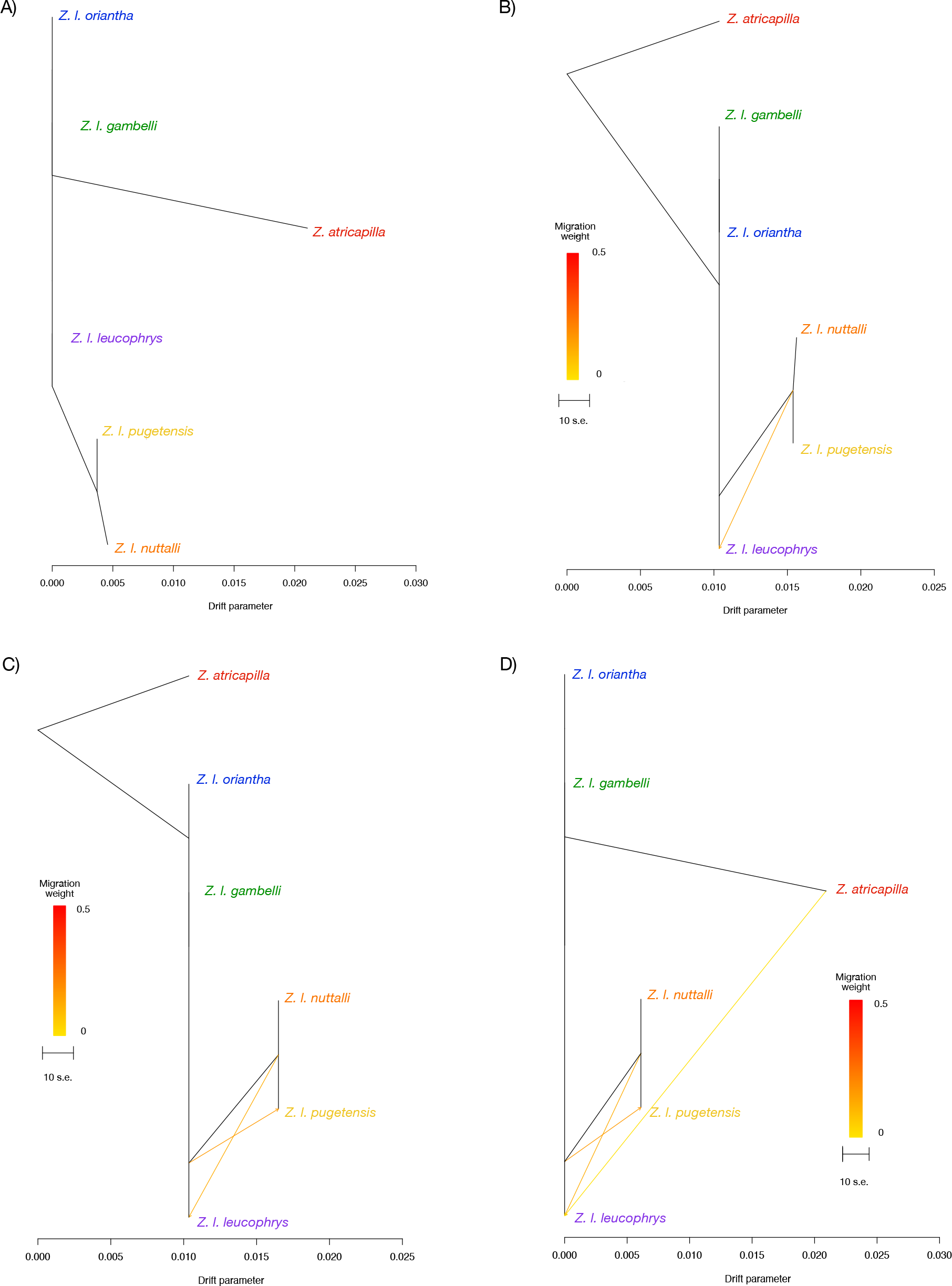
Maximum likelihood tree produced in Treemix (A), when inferring one migration event as suggested by OptM (B), when inferring two migration events (B) and three migration events (D). The bottom axis shows the drift parameter. Name labels are coloured according to species and subspecies (red=*Z. atricapilla*; orange=*Z. l. nuttallii*; yellow=*Z. l. pugetensis*; green=*Z. l. gambelii*; purple=*Z. l. leucophrys*; blue=*Z. l. oriantha*).

The ADMIXTURE analysis for K = 2 (which had the lowest cross validation error) shows *Z. atricapilla* as a different population from *Z. leucophrys*, although with some *Z. atricapilla* ancestry apparent in the GOL clade (Fig. 6A). The results for K = 3 split the three clades seen in the phylogenetic analyses (*Z. atricapilla*, NP, and GOL) with a small amount of GOL ancestry showing in the NP group, especially in *Z. l. pugetensis* individuals (Fig. 6B). For K = 4 we see some individuals within the GOL clade splitting off, and for K = 5 we see *Z. l. pugetensis* splitting off with admixed individuals in *Z. l. nuttalli* (Fig. 6C and 6D). The PCA reflected the same pattern, with the same three major groups, *Z. atricapilla*, NP, and GOL, well separated clusters (Fig. 7A). Results from the PCA on only individuals from the NP clade, shows *Z. l. nuttalli* and *Z. l. pugetensis* to be separate clusters (Fig. 7B). Similarly, when analyzing individuals from the GOL clade alone, we see Z*. l. gambelli, Z. l. oriantha*, and *Z. l. leucophrys* comprising unique clusters (although sampling is limited for *Z. l. leucophrys*; Fig. 7C). Pairwise FST between *Z. atricapilla* and all *Z. leucophrys* subspecies were high ranging from 0.27 to 0.31 (Table 2). The values were high between *Z. l. nutalli* and all other subspecies except *Z. l. pugetensis*, and were low between the subspecies within the GOL clade (Table 2). Pairwise FST values between the three major clades (*Z. atricapilla*, NP, and GOL) exceed 0.10 (Table 3).

**Fig. 6.**
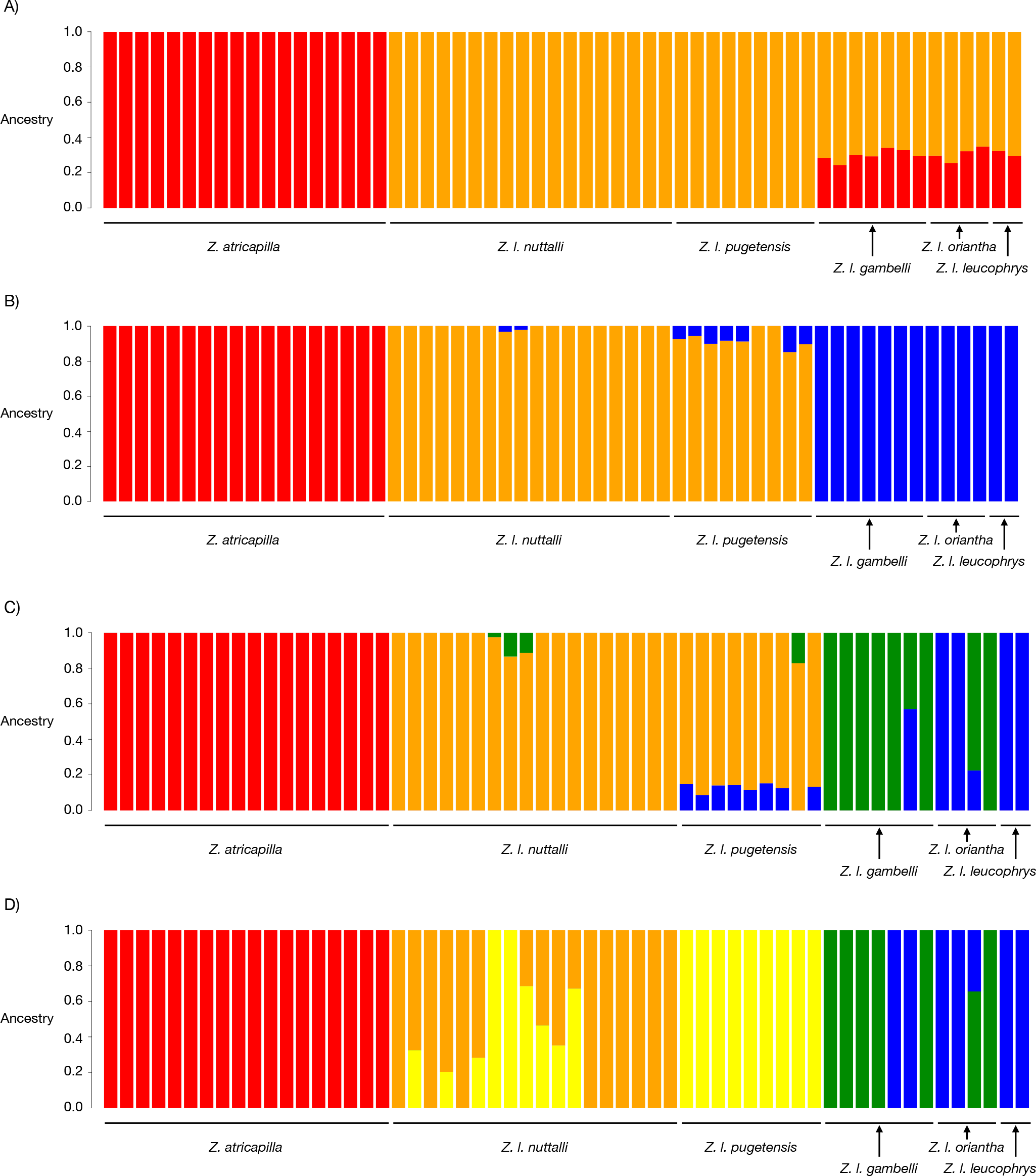
Molecular assignments of *Z. atricapilla* and *Z. leucophrys* individuals using ADMIXTURE. (A) The plot for K= 2 separating *Z. atricapilla* from *Z. leucophrys*, (B) The plot for K = 3 separating *Z. atricapilla* as one cluster*, Z. l. nuitallii* and *Z. l. pugetensis* (NP) as another, and *Z. l. gambelii, Z. l. oriantha*, and *Z. l. leucophrys* (GOL) as the third, and (C) The plot for K = 4 and (D) K = 5 showing some structuring within the NP and GOL clades.

**Fig. 7.**
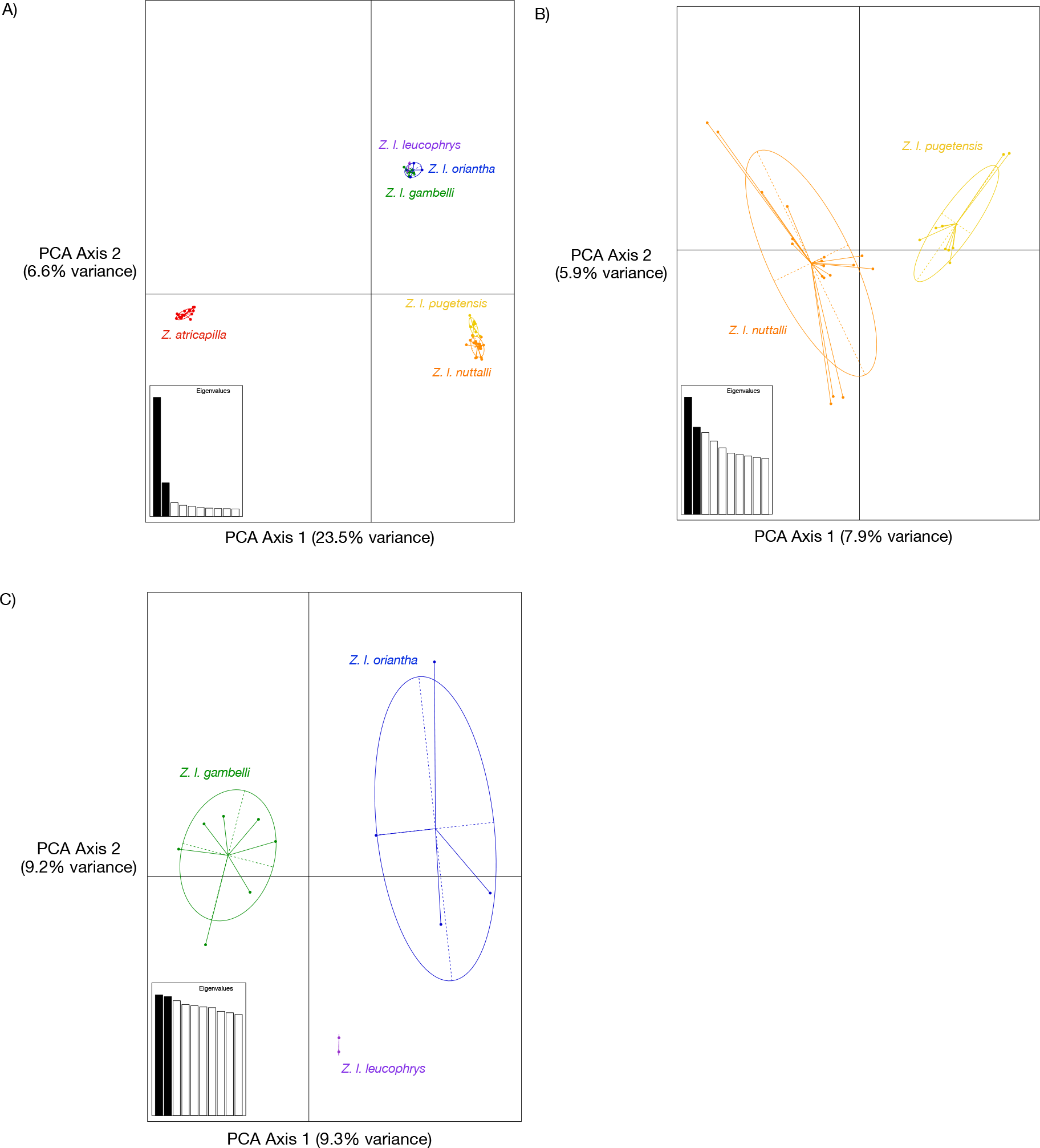
Results of a principal component analysis of *Z. atricapilla* and the five *Z. leucophrys* subspecies (A), of the NP clade (B) and the GOL clade (C). Ellipses and colours are encoded as follows: red=*Z. atricapilla*; orange=*Z. l. nuttallii*; yellow=*Z. l. pugetensis*; green=*Z. l. gambelii*; purple=*Z. l. leucophrys*; blue=*Z. l. oriantha.*

**Table 2.** Pairwise F_ST_ values from analysis of nuclear SNP data for *Z. atricapilla* and all five *Z. leucophrys* subspecies.

|  | *Z. atricapilla* | *Z. l. nuttallii* | *Z. l. pugetensis* | *Z. l. gambelii* | *Z. l. oriantha* |
| --- | --- | --- | --- | --- | --- |
| *Z. l. nuttallii* | 0.31 |  |  |  |  |
| *Z. l. pugetensis* | 0.31 | 0.04 |  |  |  |
| *Z. l. gambelii* | 0.27 | 0.13 | 0.13 |  |  |
| *Z. l. oriantha* | 0.28 | 0.13 | 0.13 | 0.01 |  |
| *Z. l. leucophrys* | 0.28 | 0.13 | 0.14 | 0.02 | 0.02 |

## 4. Discussion

### 4.1. Mitonuclear discordance in crowned sparrows

Our study is the first to investigate the phylogenetic relationships among the five *Zonotrichia* species using Next Generation Sequencing data. Contrary to inferences using mtDNA exclusively that indicated paraphyly between white-crowned and golden-crowned sparrows (Fig. 2 and 3) (Weckstein et al., 2001; McKereghan-Dares, 2008), we find *Z. atricapilla* and *Z. leucophrys* comprise well-supported, reciprocally-monophyletic sister species, with all five *Zonotrichia* species monophyletic (Fig. 4A and 4B). *Zonotrichia atricapilla* and *Z. leucophrys* comprise distinct clusters in ADMIXTURE and PCA (Fig. 6 and 7) with high FST values between *Z. atricapilla* and all *Z. leucophrys* subspecies (Tables 2 and 3). Our results suggest that *Z. atricapilla* and *Z. leucophrys* are more deeply diverged in their nDNA than in their mtDNA, and that their origins probably markedly predate the previously reported 50,000 years given the reciprocal monophyly of the species using genome wide SNP data. Consistent with Weckstein et al. (2001), our analyses imply that mitochondrial introgression resulting from hybridization is the primary cause of discordance between the two genomes for the *Z. atricapilla*-*Z. leucophrys* clade, and this strong discordance between nuclear and mitochondrial DNA generally rules out incomplete lineage sorting (Toews and Brelsford, 2012). We also found genetic structuring within *Z. leucophrys*, uncovering two reciprocally-monophyletic phylogenetic groups (Fig. 4A and 4B), which form distinct clusters in ADMIXTURE and PCAs (Fig. 6 and 7).

**Table 3.** Pairwise F_ST_ values from analysis of nuclear SNP data between the three genetic clusters uncovered in our analyses (*Z. atricapilla*, NP and GOL) – see Figure 4. NP = cluster comprising *Z. l. nutalli* and *Z. l. pugatensis.* GOL = cluster comprising *Z. l. gambelli, Z. l. oriantha*, and *Z. l. leucophrys*.

|  | *Z. atricapilla* | NP |
| --- | --- | --- |
| NP | 0.30 |  |
| GOL | 0.27 | 0.12 |

### 4.2. Causes of mitonuclear discordance

Because the vertebrate mitochondrial genome has an effective population size that is approximately one-quarter that of nDNA (Funk and Omland, 2003), we expect the rate of stochastic lineage sorting to be commensurably faster in mtDNA, and thus to exhibit differentiation more quickly in good, reproductively-isolated species (Funk and Omland, 2003). However, when hybridization and subsequent mitochondrial introgression has occurred then we can see the opposite pattern. For example, if female hybrids backcross with males of their paternal lineage over multiple generations this could result in populations where individuals bear the paternal species’ nuclear genome, and the maternal species mtDNA (Funk and Omland, 2003; see Weckstein et al., 2001).

For taxa where the fitness of heterogametic hybrids is significantly reduced, the Z (or X) chromosome is largely thought to be responsible (Haldane’s rule; Haldane, 1922; see Schilthuizen et al., 2011). This can promote speciation by preventing hybridization and fusion of incipient species (“large-Z effect”; Turelli and Orr, 1995), and thus limit opportunities for mtDNA to cross species boundaries via heterogametic hybrids (see Rheindt and Edwards, 2011). However, absolute inviability in hybrids of incipient bird species is relatively rare, and it generally takes a minimum of five million years for complete post-zygotic isolation in birds to evolve (Lijtmaer et al., 2003). As natural hybrids between these *Z. atricapilla* and *Z. leucophrys* are so rare, we have little information on their fitness in the wild.

Weckstein et al. (2001) hypothesized that, at one point in their histories, hybridization took place between *Z. leucophrys* and *Z. atricapilla* with subsequent mitochondrial introgression from one into the other. Our Treemix analysis did indicate an introgression event from *Z. atricapilla* into *Z. l. leucophrys* when adding three migration events (Fig. 5D). Under this scenario, hybridization followed by continued backcrossing of hybrids would explain the discordance between the *Zonotrichia* mitochondrial gene and species trees (Zink et al., 1991; Weckstein et al., 2001). The originally reported divergence time estimate of 50,000 years thus reflects a period of historical hybridization that essentially “reset” the mitochondrial molecular clock for the two species (see Rheindt and Edwards, 2011).

Recent evidence suggests mitochondrial introgression may be more common than previously appreciated, occurring across a wide range of taxa (Toews and Brelsford, 2012). Several processes can facilitate mtDNA introgression once hybridization has occurred. Neutral processes, such as small population sizes and genetic drift (for example during bottlenecks or founder events), or large disparities in population sizes upon secondary contact, can facilitate mitochondrial introgression (Ballard and Whitlock, 2004; Toews and Brelsford, 2012). For example, mitochondrial introgression between chipmunk species in the genus *Tamias* is thought to have occurred due to low population densities during climate induced range shifts (Good et al., 2008), and differential population sizes may have promoted mitochondrial introgression in *Neodriprion* sawflies (Linnen and Farrell, 2007).

Mitochondrial introgression can also be caused by selection (i.e. adaptive introgression; Toews and Brelsford, 2012). Increasing evidence highlights the importance of selection on the mitochondrial genome, contradicting the long-held assumption that much of the variation in mtDNA can be regarded as neutral (Ballard and Whitlock, 2004; Toews and Brelsford, 2012). For example, selection on the mitochondrial genotype (or perhaps other cytoplasmic factors) is likely to have caused the introgression of mtDNA between *Drosophila teissieri* and the *D. yakuba/D. santomea* species pair (Bachtrog et al., 2006). Similarly, a mitochondrial selective sweep is thought to have caused mitonuclear discordance in yellowhammers (*Emberiza citrinella*) and pine buntings (*E. leucocephalos*; Irwin et al., 2009). Examples of adaptive introgression often invoke thermal adaptation as an explanation, due to the key role of mtDNA in efficient energy production and metabolism (Ballard and Whitlock, 2004; Lamb et al., 2018). For example, mitochondrial introgression between *Lepus* hare species (Melo-Ferreira et al., 2005), and between arctic char and lake trout (*Salvelinus alpinus* and *S. namaycush*; Wilson and Bernatchez, 1998), are potentially due to temperature adaptation. Mitochondrial variation in multiple Australian bird species is associated with historical climatic variation (Lamb et al., 2018).

It is unclear whether the mitochondrial introgression between *Z. atricapilla* and *Z. leucophrys* was caused by neutral or selective processes, or a combination thereof. The Pleistocene epoch was characterized by repeated cycles of global cooling and warming, with the most rapid oscillations occurring from 60,000 to 25,000 years before present (Adams, 1997). In the northern hemisphere, colder periods resulted in a series of extensive glacial advances. At their maximum extent Arctic ice sheets covered most of present-day Canada (Adams, 1997). Cooling also compressed and pushed temperate and tropical climates towards the equator, and promoted the expansion of desert habitats (Hewitt, 2004).

While it is only informed speculation, during the Pleistocene populations of *Z. atricapilla* and *Z. leucophrys* could have been forced into common refugia by ice sheets and shifting vegetation, leading some sexually mature adults to seek out heterospecific mates. If the newly-introgressed mitochondrial haplotype conferred favorable metabolic or physiological traits, it could have become fixed throughout populations of both species due to adaptive introgression. Several rounds of bottlenecking due to climate change might have occurred during this period and similarly caused the fixation of one haplotype in both species, or acted in combination with selection to augment this process.

If effective population sizes were small, and the young species were defined by relatively few variable sites, the re-assortment of nuclear loci could have occurred quickly to erase traces of hybridization events. Premating isolation mechanisms may have also been well-developed to prevent the fusion of the species. Behavioural reproductive isolation through conspecific song recognition is well-documented in *Zonotrichia* species, with juveniles predisposed to learn their own species’ songs (e.g. Shizuka, 2014). The distinct coloration of their crowns might have further distinguished them, but the importance of visual signaling in species recognition or mate selection remains to be explored in crowned sparrows.

The time required for biological speciation is taxon-dependent, with some species able to interbreed even after more than 20 million years of divergence (Lijtmaer et al., 2003; Coyne and Orr, 2004). Given that little gene flow is needed for mtDNA to move between species successfully (Takahata and Slatkin, 1984), and complete reproductive isolation can take millions of years to develop, mitochondrial introgression could be far more common in birds than previously realized. Literature surveys suggest that 14-16% of bird species exhibit mitochondrial paraphyly, and it is estimated that at least 5.7% of these cases are due to introgression (Funk and Omland, 2003; McKay and Zink, 2010).

### 4.3. Implications of mitochondrial introgression

An increasing number of studies are revealing the importance of selection on the mitochondrial genome (Toews and Brelsford, 2012). Despite this, the drivers of mitochondrial introgression are not well-studied in natural systems. Future work should investigate the functional differences caused by different mitochondrial haplotypes, as was done in yellow-rumped warbler species (*Setophaga* spp.; Toews et al., 2014). In addition, the co-adaptation of mitochondrial and nuclear genes is of key importance for fitness (Hill, 2017). Cases such as the *Zonotrichia* sparrows, where one mitochondrial haplotype was completely replaced by another, indicate that in some instances the cytonuclear interactions are weaker than the strength of selection on the mtDNA (Bachtrog et al., 2006).

Regardless of whether mitochondrial introgression is caused by neutral or selective processes, there are implications in the use of mtDNA alone for inferring species histories. For example, the use of both nuclear and mitochondrial DNA in Asian box turtles (*Cuora* spp.) allowed the recognition of monophyletic taxon, which had been masked by complicated patterns of mitochondrial introgression, with strong implications for the choice of individuals to use in captive breeding programs (Spinks and Shaffer, 2007). In addition, there are obvious implications for taxonomy, with introgression confounding phylogenetic inference of species (e.g. crotaphytid lizards; McGuire et al., 2007), or masking hidden genetic structuring as we have found both between *Z. atricapilla* and *Z. leucophyrys*, and among *Z. leucophrys* subspecies.

### 4.4. Relationships between the five Zonotrichia species and the Z. leucophrys subspecies

In both the mitochondrial and nuclear phylogenetic reconstructions (Fig. 3 and 4), *Z. capensis* is the most distinct and sister to all other *Zonotrichia* species. However, we see different patterns between the nuclear and mitochondrial phylogenies with *Z. querula* and *Z. albicollis* which are recovered as sister species in the mitochondrial DNA tree (Fig. 3), but *Z. albicollis* is basal to *Z. querula* sister in nuclear DNA (Fig. 4). To resolve the relationship, it may be necessary to increase the sampling of these two species to increase the power of the analyses.

We found well-supported monophyletic clades within *Z. leucophrys* that divide the subspecies into two groups (GOL and NP; Fig. 4, 6, and 7). Further, we found support for genetic differences between *Z. l. nuttallii* and *Z. l. pugetensis* (Fig. 4A, 4B, and 7B), which is consistent with the differentiation found by Lipshutz et al. (2017), although we do not find them to be completely reciprocally monophyletic with one *Z. l. pugetensis* individual recovered within the *Z. l. nuttallii* clade (Fig. 4A and 4B). We likely find stronger structuring and less evidence of hybridization between *Z. l. nuttallii* and *Z. l. pugetensis* in our study because we did not use individuals from within or near to the contact zone (Fig. 1). Within the GOL clade, however, the subspecies are not recovered as distinct (Fig. 4A and 4B) with low FST values among three subspecies (Table 2). We do find the that PCA separated them (Fig. 7C) suggesting some weak differentiation.

## 5. Conclusions

Our findings support previous assertions of historical mitochondrial introgression between the two sister species, and further emphasize the need to re-evaluate estimates of age of divergence. For example, future studies with more data could re-estimate divergence times using the relative branch lengths within the *Zonotrichia* phylogeny, as the age of other *Zonotrichia* species (e.g. *Z. capensis*) are better established (Lougheed et al., 2013). More work should also focus on better delineating the genealogical patterns of the *Z. leucophrys* subgroups. These provide an important opportunity to study different stages of the speciation process *in situ*. Finally, we show a clear need to incorporate both nuclear and mitochondrial genomes when investigating avian evolutionary histories. For example, McKay and Zink (2010) found that in 21.3% of cases of paraphyly in birds, introgression could not be reliably distinguished from incomplete lineage sorting (McKay and Zink, 2010). Mitochondrial DNA is effectively a single locus and lacks the resolution needed to discern close relationships and cannot on its own reveal signatures of hybridization. Genome-wide nDNA sequencing is more powerful, provides better resolution, and allows us to make stronger inferences about the histories of many taxa. Ultimately cases of mitonuclear discordance tell us something important about both the history of species (such as *Zonotrichia* sparrows) and about mitochondrial evolution. We must consider mtDNA as simply a ‘tool’ for phylogeography (Ballard and Whitlock, 2004).

## 6. Author statement

RST conducted analyses and contributed to the writing of the manuscript; ACB conducted lab work and contributed to the analysis and writing of the manuscript; NAC contributed to the writing of the manuscript; FB contributed to the writing of the manuscript and assisted with procuring samples, RC-C conducted lab work and assisted with writing of the manuscript. SCL conceived the study, did some phylogenetic analysis, and contributed to the writing of the manuscript.

## Acknowledgements

We thank John Wingfield (University of California, Davis); Dai Shizuka (University of Nebraska); the Burke Museum of Natural History and Culture; the Royal Alberta Museum; the Royal Ontario Museum; the University of Alaska Museum; Donna Maney (Emory University); Beth MacDougall-Shackleton (Western University); Paul Martin (Queen’s University); and the Louisiana State University Museum of Natural History for contributing tissue samples. Brian Boyle (Genomic Analysis Platform, Laval University) and Génome Québec Innovation Centre performed the library preparation and Illumina sequencing, respectively. This research was part of two undergraduate theses (ACB and KD) at Queen’s University. This work was supported by a Natural Sciences and Engineering Research Council of Canada (NSERC) Discovery Grant to S.C.L., the Baillie Family Chair in Conservation Biology (SCL), a NSERC Undergraduate Student Research Award to ACB, a NSERC Alexander Graham Bell Canada Graduate Scholarship to NAC, and an Ontario Trillium Scholarship to RST.

