## Supplementary Fig. 1 for "Cytonuclear discordance in the crowned-sparrows, *Zonotrichia atricapilla* and *Zonotrichia leucophrys*: a mitochondrial selective sweep?"

**Fig. S1.** The locations of sampling sites of *Z. atricapilla, Z. leucophrys, Z. albicollis, Z. capensis,* and *Z. querula* individuals   The background colors represent the breeding ranges, pink for *Z. atricapilla* and purple for *Z. leucophrys* with deep pink representing range overlap. More sampling details are provided in Table 1.

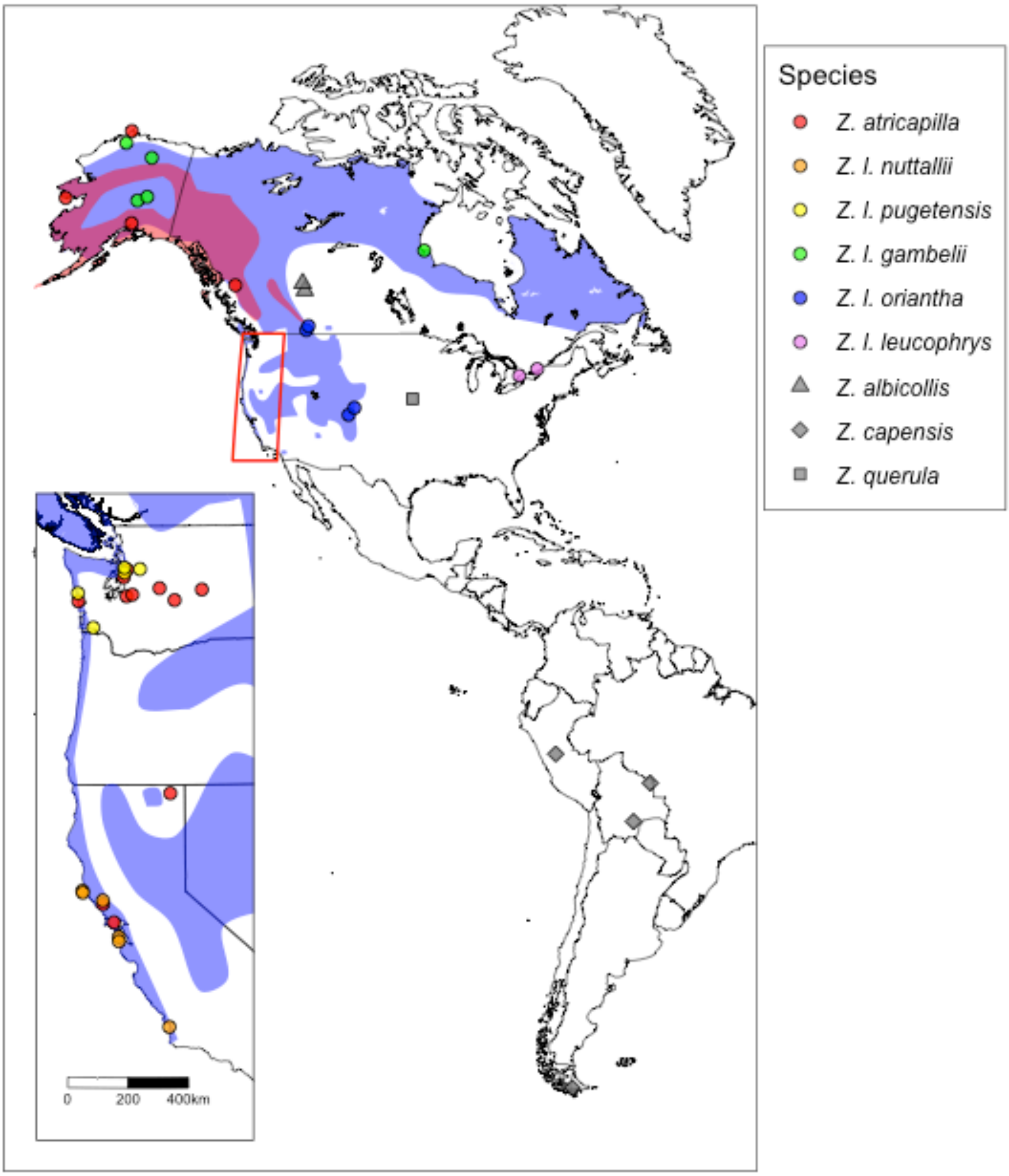
