## Supplementary Fig. 2 for "Cytonuclear discordance in the crowned-sparrows, *Zonotrichia atricapilla* and *Zonotrichia leucophrys*: a mitochondrial selective sweep?"

**Fig. S2.** Bayesian mitochondrial DNA phylogenetic analyses from all five *Zonotrichia* subspecies.

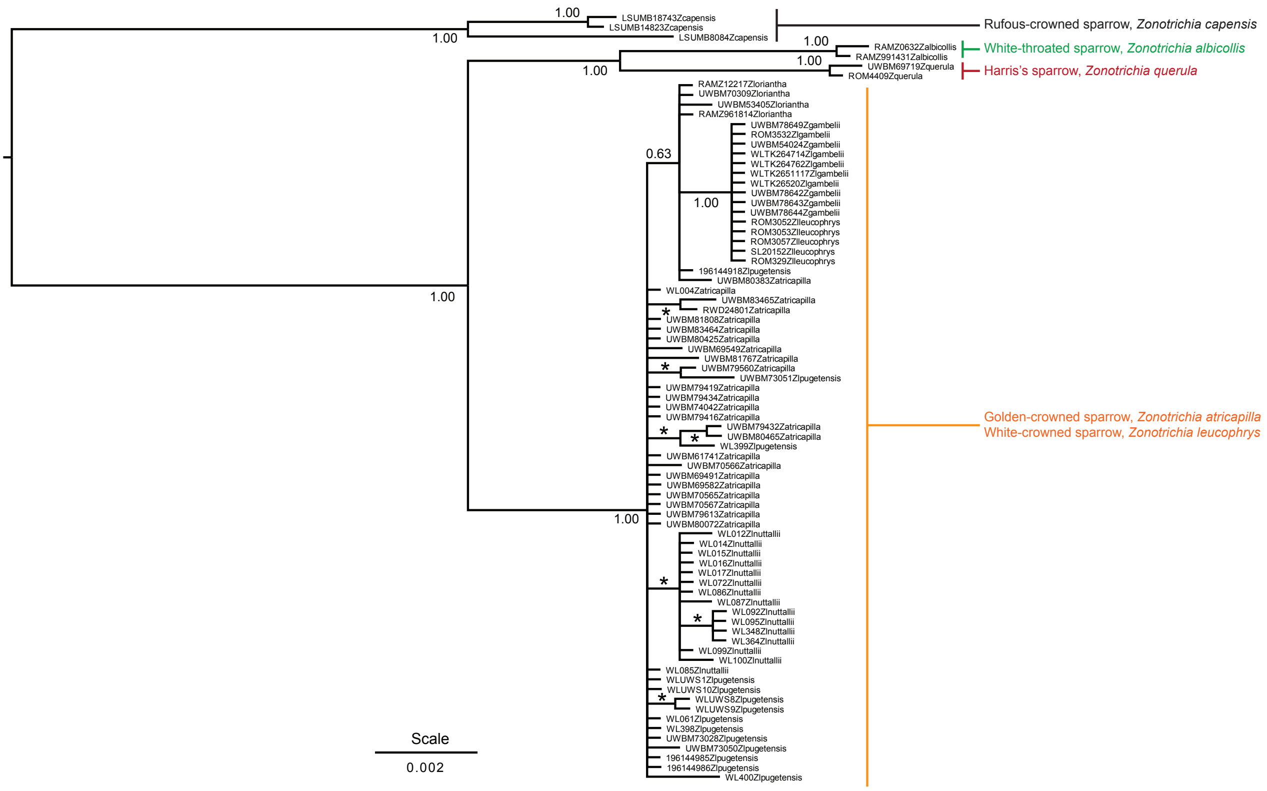
