## Supplementary Fig. 3 for "Cytonuclear discordance in the crowned-sparrows, *Zonotrichia atricapilla* and *Zonotrichia leucophrys*: a mitochondrial selective sweep?"

**Fig. S3.** OptM results showing the variance explained by adding migration events in Treemix (top) and the delta m statistic for each number of migration events (bottom).

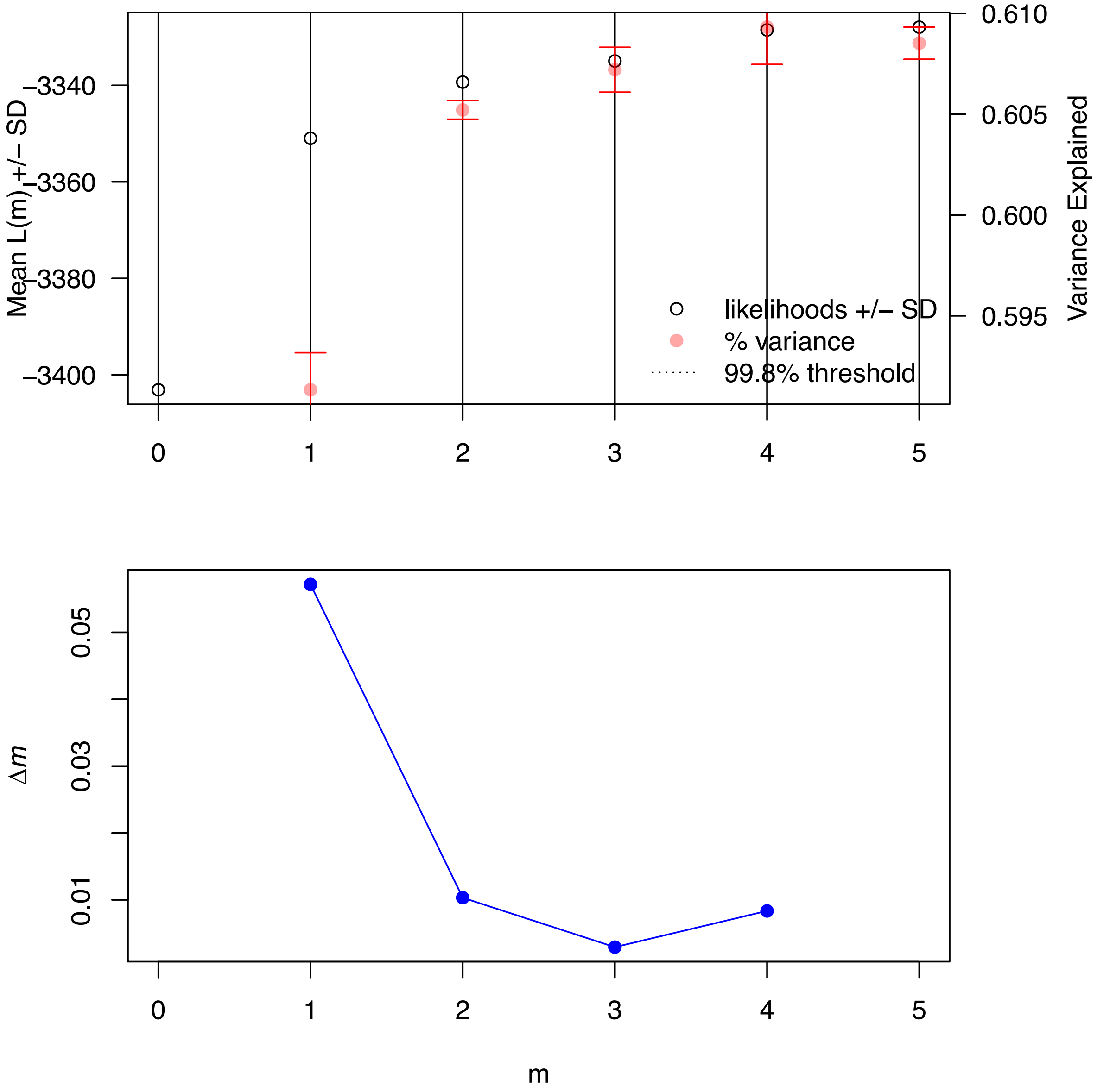
